# Neural tube patterning: from a minimal model for rostrocaudal patterning towards an integrated 3D model

**DOI:** 10.1101/2020.10.02.323535

**Authors:** Max Brambach, Ariane Ernst, Sara Nolbrant, Janelle Drouin-Ouellet, Agnete Kirkeby, Malin Parmar, Victor Olariu

## Abstract

The rostrocaudal patterning of the neural tube is a key event in early brain development. This process is mainly driven by a gradient of WNT, which defines the fate of the present neural progenitor cells in a dose dependent matter and leads to a subdivision of the tube into forebrain, midbrain and hindbrain. Although this process is extensively studied experimentally both *in vivo* and *in vitro*, an integrated view of the responsible genetic circuitry is currently lacking. In this work, we present a minimal gene regulatory model for rostrocaudal neural tube patterning. The model's nodes and architecture are determined in a data driven way, leading to a tristable configuration of mutually repressing brain regions. Analysis of the parameter sensitivity and simulations of knockdown and overexpression cases show that repression of hindbrain fate is a promising strategy for the improvement of current protocols for the generation of dopaminergic neurons *in vitro*. Furthermore, we combine the model with an existing model for dorsoventral neural tube patterning, to test its capabilities in an *in vivo* setting, by predicting the steady state pattern of a realistic three-dimensional neural tube. This reveals that the rostrocaudal pattern stacks dorsoventrally in the caudal half of the neural tube. Finally, we simulate morphogen secretion overexpression, which highlights the sensitivity of neural tube patterning to the morphogen levels.

## Introduction

In Parkinson’s disease (PD), the dopamine-producing (dopaminergic) neurons (DA) of the substantia nigra of the brain undergo pathological deterioration. The resulting deficiency in striatal dopamine leads to symptoms such as tremor, bradykinesia and rigidity that can currently not be relieved sufficiently by pharmacological treatment (1,2). Today, transplantation therapy is the most promising approach to achieving PD recovery (3); several studies and clinical trials have demonstrated restoration of dopaminergic activity and partial symptom relief for up to 18 years post-transplantation (2,4,5). The conventional cell source for these transplants, human fetal ventral mesencephalic tissue – which contains a high concentration of dopaminergic neuroblasts (4,6) – is naturally scarce, ethically controversial and suffers from highly variable results (6).

Recently, it has been suggested that DA progenitor generation by differentiation of human pluripotent stem cells (hPSCs) presents an alternative approach. Studies in animal models have shown that hPSC-derived DA progenitors are equivalent to fetal cells in terms of subtype specific marker expression, controlled dopamine release and functional PD symptom relief (3,7–10). To date, various protocols have been developed to efficiently derive DA progenitors from human embryonic stem cells; many of them focusing on tuning a combination of WNT, SHH and FGF8 with great success (11–16). These protocols are inspired by the mechanisms and actors found in the development of the neural tube *in vivo*. Therefore, a better understanding of correct and reliable DA differentiation *in vivo* would not only shine more light on the development of the early brain, but also contribute to improving current *in vitro* protocols for midbrain DA neuron generation.

*In vivo* neural patterning is a complicated and concerted process that relies on the three-dimensional diffusion of various morphogen signals in an uneven geometry. In humans, the current understanding of this process can be summarised as follows. By the end of the fourth week of embryonic development the progenitor of the embryo’s brain, the neural tube, is patterned along the rostrocaudal axis into three distinct regions that develop into forebrain, midbrain and hindbrain (17). In addition, patterning occurs along the dorsoventral axis, setting up numerous progenitor cell types such as motor neurons and interneurons (18). The formation and subdivision of the neural tube is the result of extracellular morphogens that are released in specific areas of the tube, establishing concentration gradients. The cells of the neural tube are able to interpret the local concentration of these gradients and take fate decisions accordingly. Dorsoventral patterning is mainly controlled by opposing signalling gradients of WNT/BMP from the roof plate, and SHH from floor plate cells (19) and the zona limitans intrathalamica (20). Since gene regulatory networks have been shown to control pattern formation in multiple tissues (21–24), the developing vertebrate neural tube pattern might be the response of transcriptional circuitry inside neural progenitor cells to morphogen gradients.

Neural tube rostrocaudal patterning is mainly governed by WNT-signalling, emerging from the isthmic organizer (25). It has been shown in vitro that controlling the gradient of the WNT-signalling, using GSK3 inhibitors either in different cell cultures (11) or in microfluidic devices (26), acting on differentiating human embryonic stem cells can lead to progressive caudalisation from forebrain to hindbrain. However, the gene regulatory network acting downstream of WNT-signalling that regulate this patterning processes has not been elucidated.

In this study, a minimal gene regulatory network model for rostrocaudal neural tube patterning is proposed. The transcription circuit topology was determined in an unbiased way and the model parameters were optimised using gene expression data from a study on hPSCs, which have been cultured in conditions with varied levels of WNT-signalling. Subsequently, model parameter sensitivity analysis revealed the key interactions that drive the patterning process. Moreover, knockdown and overexpression model simulations gave insight into the regulation of the patterning and enabled the prediction of more efficient protocols for the generation of DA neurons *in vitro*. Finally, our optimised model was combined with the model controlling dorsoventral patterning proposed in (27) to simulate patterning of the neural tube in a realistic, three-dimensional geometry. Our findings represent a step forward towards a complete understanding of neural patterning and towards an optimal DA progenitor derivation protocol. Furthermore, recent advances in PD research incorporating organoids (28,29) make it necessary to not only investigate morphogen and gene interactions but also the effects of tissue geometry on neural patterning.

## Results

### Gene expression of GSK3i treated cells clusteres three-fold

As a first step towards a minimal gene regulatory network for rostrocaudal neural tube patterning, we aimed to identify the essential genes involved in this process. To this end, we used experimental data consisting of the expression levels of a selection of genes recorded in hPSCs for varying concentrations of the GSK3 inhibitor CHIR99021 (CT). This dataset has been published in Kirkeby et. al (11). GSK3 is a negatively regulated target of canonical WNT-signalling, i.e. low GSK3 levels emulate strong WNT activity and vice versa. To identify the genes that correspond to the patterned brain regions, we calculated the correlation between the gene expression across the varied CT concentrations. For that, we used a hierarchical agglomerative clustering approach, resulting in a correlation cluster map (30) (Figure 1 A). From this analysis, we identified the three major clusters with positive correlation. Examining the expression levels of these three gene clusters, three distinct regions of gene expression are found for varied levels of CT (Figure 1 B). For the gene regulatory network model parameter and topology optimisation we selected one representative gene for each brain region. For the forebrain (FB) region we used forkhead box G1 (FoxG1), since it has been shown to be involved in telencephalon fate (31–33). The midbrain (MB) region was represented by engrailed 1 (EN1), due to its key role in DA neuron specification (34,35). For the hindbrain (HB) region we chose HOXA2, as it is reported to be a central regulator in hindbrain fate (36,37). The selection of one representative gene per brain region enabled us to construct a minimal gene regulatory network to capture the dynamics of neural tube patterning. However, it is important to note that this does not necessarily imply that the selected genes are interacting directly or are primarily regulating cell fate specification. Therefore, in the following results, the labels FB, MB and HB are used, when referring to the genes.

**Figure 1.**
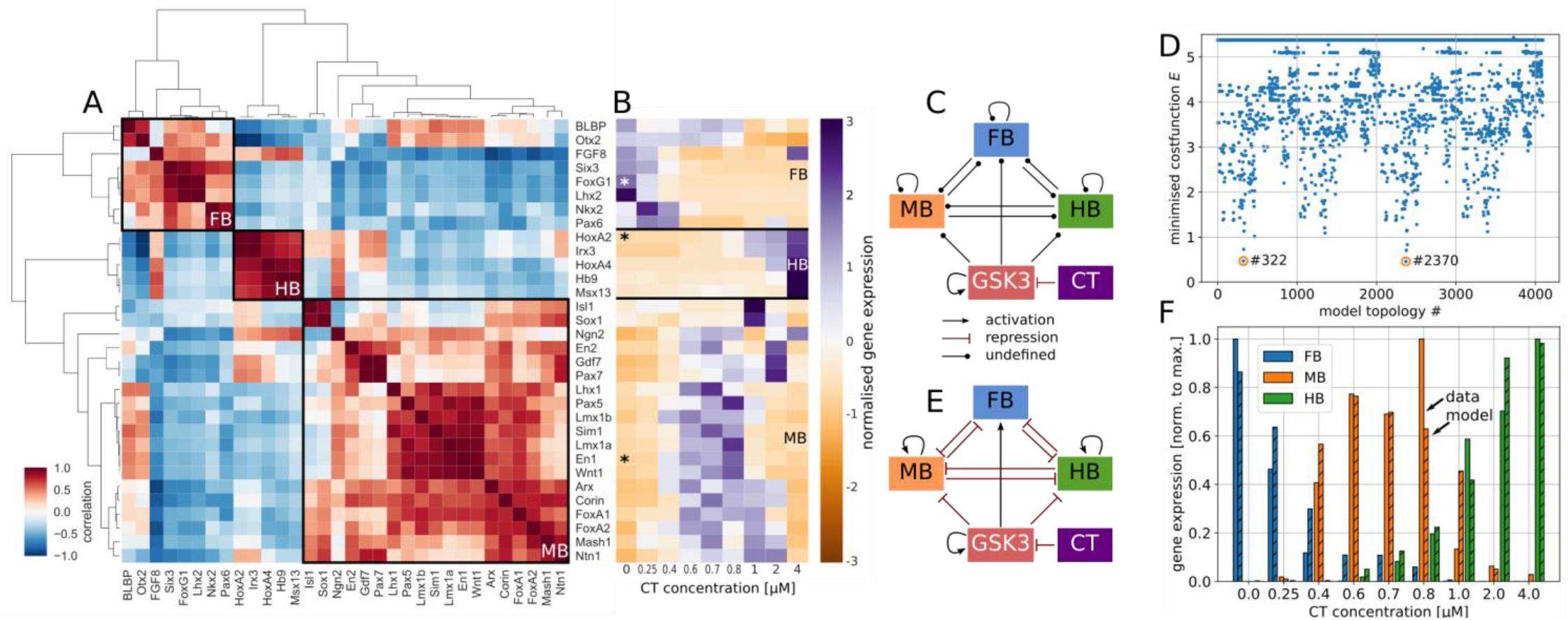
Experimental data from *in vitro* neural patterning experiments. A: Correlation cluster map of the different brain region specific genes across CT concentrations. Three clusters were identified, which correspond to the three brain regions. B: Normalised gene expression of FB, MB and HB specific genes for different concentrations of the GSK3 inhibitor CT. Genes marked with asterisk are used to represent the brain regions in the following results. C: Model network with all possible interactions between the brain region nodes and their reaction dependency on GSK3 expression. CT controls GSK3, which is self-activating to allow for steady-state GSK3 expression. During the topology selection, all undefined interactions were either activation or repression. D: Minimal optimised cost function value for all possible network topologies. Two best topologies with similar and overall lowest cost functions were identified. E: Best model as determined by the topology selection. The three brain regions formed a tristable switch, controlled by the level of GSK3, which is in turn controlled by CT concentration. Note that GSK3, MB and HB are self-activating to allow for respective constant steady state gene expression. Degradation of GSK3, FB, MB, HB not illustrated. F: Gene expression levels of the three brain region specific genes given by the experimental data (plain bars) and the model (striped bars) for varied concentrations of CT.

### Unbiased topology selection identifies a tristable switch configuration

With the key players of the network identified, the next step towards a minimal gene regulatory network was the determination of their interactions i.e. the connection between the nodes. These nodes are FB, MB, HB, GSK3 and CT (Figure 1 C). We determined the interaction between these nodes and the resulting network topology in an unbiased way by optimising all possible configurations towards the experimental data. The only two interactions that were pre-defined are the repression of GSK3 by CT, which results in the necessity of GSK3 self-activation to allow for steady state expression of GSK3. The remaining interactions can be either activation or repression, which makes for 4096 possible network configurations. The configurations that yield the lowest cost function *E_d_* (see Methods) after parameter optimisation were considered as the winning configurations (Figure 1 D); two topologies satisfy this criterion. These two configurations only differ in the self-interaction of FB, which is activation in one case and repression in the other. However, the corresponding rate constant *c*_0_ (see Methods) had a significantly smaller value compared to the other rate constants and therefore we concluded that, for the purpose of this minimal model, FB does not self-interact.

The model resulting from the unbiased topology selection is shown in Figure 1 E. In this model, the three brain regions are mutually repressive and hence form a tristable switch. MB and HB fate are downregulated by GSK3, whereas FB fate is positively regulated. Also, FB does not self-interact and therefore directly follows GSK3 expression when the latter is expressed at high levels. For lower levels of GSK3, FB is no longer strongly expressed and can be fully repressed by the presence of MB or HB. This reduces the network to a bistable switch between MB and HB fate for lower GSK3 levels, which is controlled by their mutual repression strength and the repression through GSK3. In fact, the corresponding parameters were determined such that HB is repressing MB much stronger than vice-versa. However, GSK3’s repression on HB is even stronger compared to its repression of MB. It follows that for low GSK3 levels HB will dominate over MB and vice versa for higher GSK3 levels.

Figure 1 F shows the brain regions’ expression levels computed by using the model together with the experimental data used for the parameter optimisation. For most CT concentrations, model and data show high correlation. A significant deviation between model and data only occurred for the CT concentrations 0.8 μM and 1.0 μM. However, it is important to note that the overall structure of the brain regions’ response to varied CT concentration in this minimal model is close to the experimental data.

### Sensitivity analysis of model parameters reveals key interactions

Next, we examined the sensitivity of the model output to changes of the model parameters to identify the key interactions. *In vitro* data were used as reference and consequently the results are given in terms of the cost function *E_d_*, which was defined as the mean squared distance between model and data. Figure 2 A, B show that the most sensitive rate constants are *c*_4_, *c*_8_ and *c*_12_, which correspond to the self-activation of MB, HB and GSK3. A similar sensitivity pattern was found in the Hill coefficients *n*_*i*_, for which also *n*_4_, *n*_8_, and *n*_12_were the most sensitive parameters, illustrating cooperativity and ultrasensitivity for MB, HB and GSK3. The most sensitive degradation rates were also the ones corresponding to the half-lives of MB, HB and GSK3 respectively (*δ*_2_, *δ*_3_ and *δ*_4_). This implied that the most sensitive part of the model was the balance between self-activation and degradation of MB, HB and GSK3 expression and the resulting steady state gene expression level. This was to be expected, since those expression levels are controlling which node of the tristable switch is active. FB does not self-interact, and its expression is therefore strongly correlated to the expression of GSK3 at high GSK3 levels. Also, high GSK3 levels lead to repression of MB and HB, whereas at low GSK3 levels MB or HB repress the expression of FB, explaining the low sensitivity of the model to changes of the FB degradation rate *δ*_1_. The sensitivity of the repressions between FB and HB (*c*_2,10_, *n*_2,10_) was low, because either (at high GSK3 levels) HB was repressed by GSK3 or (at low GSK3 levels) FB was turned off. In both cases the sensitivities of the rate constants and Hill coefficients were low because the corresponding concentrations were close to zero. Another weak sensitivity was found in the repression of HB by MB (*c*_9_, *n*_9_), implying that the HB level was more dependent on its self-balance and repression by GSK3 than on repression by MB. Figure 2 C shows the mean cost function change when all parameters were varied. We observed that the model was not sensitive to small variations of the model parameters. The mean cost function change for the variation of the whole parameter set of ± 10 % was similar to the maximum cost function change for the variation of an individual parameter. This showed that the model is robust under the variation of parameters.

**Figure 2.**
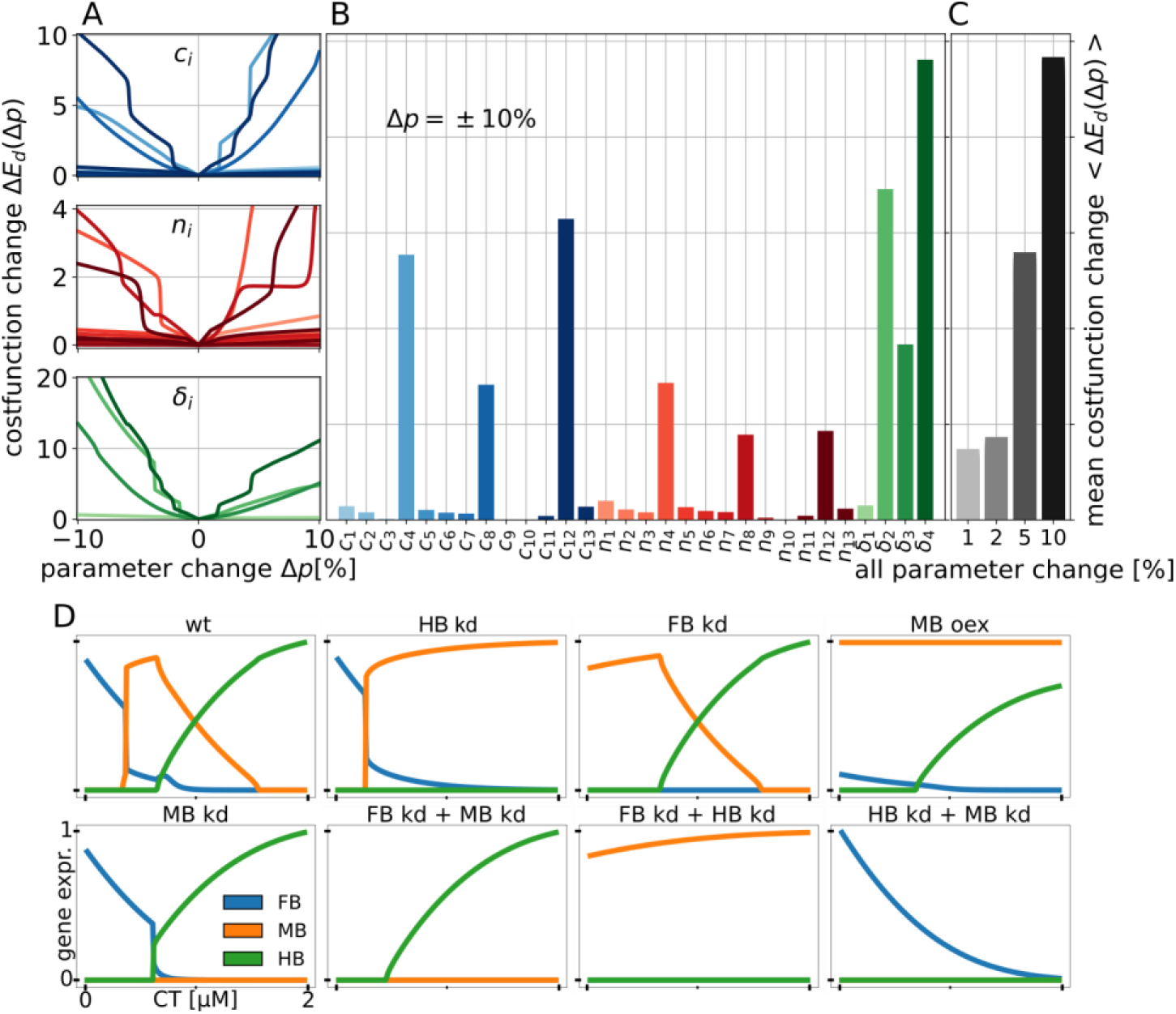
Sensitivity analysis of the model parameters and kd / oex predictions. A: Fold-change of the cost function **Δ*E_d_***(**Δ*p***), with **Δ*p*** being a parameter set with one parameter varied between 0 and 10 % relative to the optimised parameter set. B: Mean cost function change 〈**ΔE**(**Δ*p***) 〉 across **Δ*p*** = ± **10** % relative to the optimised value. C: 〈**Δ*E_d_***(**Δ*p***)〉 for all parameters varied ± different relative amounts to the optimised parameter set. D: HB kd showed strong MB expression for CT > 0.2 μM. FB kd lead to strong MB expression for CT < 0.3 μM and wt behaviour for higher CT concentrations. MB oex suppressed FB expression and lead to mixed MB/HB expression for higher CT concentrations. MB kd gave insight into the interaction between HB and FB and the double kd’s highlight the reaction of the brain regions’ gene expressions to the CT concentration. Axes are the same for all panels, gene expression is in arbitrary units between 0 and 1.

The sensitivity analysis shows that the key parameters of our model are the self-interactions of MB, HB and GSK3, since they not only control their own expression level, but also which node of the tristable switch is active. Furthermore, the general dynamics of the model seem to be that FB is proportional to GSK3, MB and HB are anti-proportional to GSK3 and the extent of the MB peak is controlled by the HB expression level. Therefore, we can formulate the hypothesis that knockdown of FB would not have a strong effect on the extend of the MB domain. However, knockdown of HB would remove the repression of MB for low GSK3 levels and would therefore lead to an extended MB region that includes the former HB region.

### Knockdown and overexpression simulations show that only HB knockdown increases the MB domain

To test this hypothesis and to gain more insight into the transcriptional regulation of this patterning, we simulated knockdown (kd) and over-expression (oex) cases (Figure 2 D). HB kd did not influence the interaction between FB and MB and lead to strong MB expression for CT > 0.2 μM, resulting in a more than five-fold larger CT window for MB fate. Knockdown of FB similarly did not influence the interaction between MB and HB and led to strong MB expression for CT < 0.3 μM. However, this only doubled the extent of the CT window for MB fate. Interestingly, when overexpressing MB, FB expression was lost, but the HB expression pattern was similar to wildtype. This gave rise to mixed MB-HB expression for CT > 0.4 μM and resulted in a CT window for HB fate similar to the FB kd case. Knockdown of MB revealed that FB and HB have the same mutually exclusive interaction as FB and MB in the wildtype case. This produced a slightly larger FB domain, whereas the HB domain was almost unaffected in extent or shape. The double knockdown simulations of FB and MB showed that HB expression requires a threshold CT concentration of CT > 0.2 μM and reacts positively to increasing CT past that threshold. Knockdown of MB and HB illustrated that FB expression is negatively regulated by CT. Interestingly, the knockdown of FB and HB showed that MB reacts only weakly to CT at low levels. This implies that MB is only weakly regulated by CT itself.

These results confirmed the findings from the parameter sensitivity analysis and suggest that protocols for the generation of DA MB neurons could be made more robust and independent of precise CT concentration by repressing HB fate. This would significantly increase the window of CT concentration that leads to MB fate, while the repression of FB fate or the promotion of MB fate via overexpression would not lead to the same result.

### Application of model to realistic geometry revealed dorsoventral stacking of rostrocaudal pattern

To further test the potential of the minimal rostrocaudal model, we merged it with the model of Balaskas et al. for dorsoventral patterning (27) and used the combined model to simulate the steady state pattern of the human neural tube *in vivo*. The combined model is shown in Figure 3 A. WNT signalling controls both dorsoventral and rostrocaudal patterning and therefore, the two model branches are linked through the WNT node. In the rostrocaudal network branch, a buffer node (U) translates the WNT signal into the GSK3-inhibition as achieved by CT in the *in vitro* case. On the dorsoventral network branch, Gli expression is repressed by WNT signalling and activated by SHH activity (19). Gli expression acts as selector for dorsoventral fate – like GSK3 in the rostrocaudal case – by activating the expression of Nkx2.2 (ventral fate, N) and Olig2 (central fate, O), which are mutually repressive. Both N and O repress the expression of Pax6 (dorsal fate, P), resulting in a reduced tristable switch network motif.

**Figure 3.**
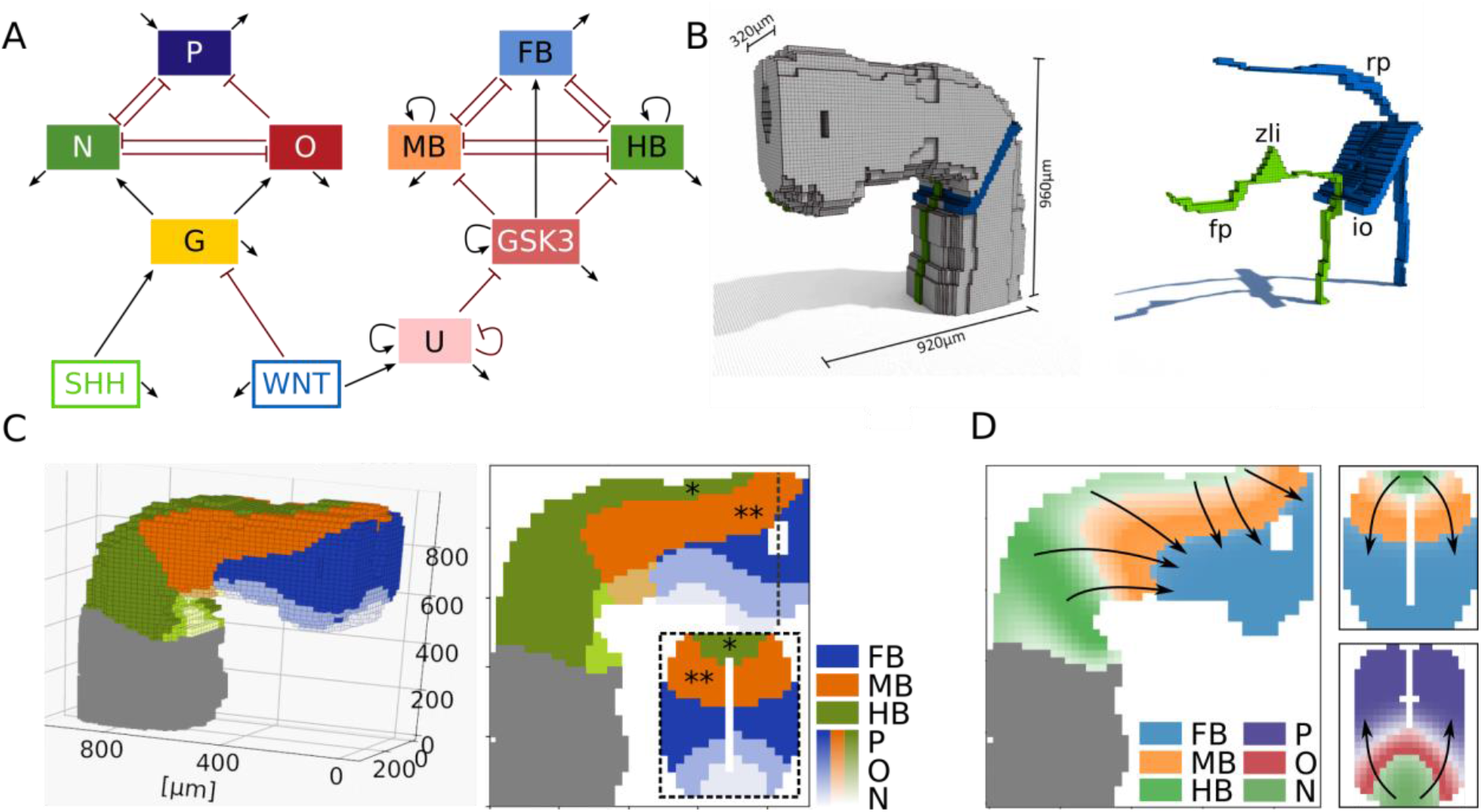
In vivo patterning of the neural tube. A: Complete model for neural tube patterning. Left branch: dorsoventral model by *Balaskas et al.,* controlled by antagonistic action of SHH and WNT. G: Gli, P: Pax6, O: Olig2, N: Nkx2.2. Right branch: Proposed model for rostrocaudal patterning governed by WNT action. B: Three-dimensional model of neural tube geometry and the morphogen secretion sources. SHH (green): floor plate (fp) and zona limitans (zli); WNT (blue): roof plate (rp) and isthmic organiser (io). C: Steady-state distribution of the patterned regions after applying the complete model. The geometry of the tube caused dorsoventral stacking of the rostrocaudal gene domains (* HB stacking, ** MB stacking). D: Rostrocaudal and dorsoventral steady-state pattern. Arrows indicate the direction of pattern establishment i.e. the main direction of morphogen action. Greyed areas in C and D are only shown for orientation and are not considered for pattern establishment.

To explore effects of the curved tube geometry and morphogen secretion sites on patterning, we set up a spatial simulation of the network on a realistic three-dimensional model of the rostral neural tube (38) (Figure 3 B). The WNT signal originates from the roof plate (rp) and the isthmic organiser (io) and SHH is secreted from the floor plate (fp) and the zona limitans intrathalamic (zli) (20). The diffusion dynamics of both morphogens were estimated based on their respective protein structure (see Methods).

The calculated steady state pattern (Figure 3 C) agreed with previous studies for the dorsoventral branch (27) and also yielded the anticipated rostrocaudal pattern in the ventral half of the neural tube. Interestingly, in the dorsal half of the neural tube, the additional WNT signal from the roof plate lead to a stacking of the rostrocaudal brain regions in dorsoventral direction (* HB stacking, ** MB stacking in Figure 3 C, D), which resulted in an L shaped MB and HB region in a sagittal cross section. A frontal cross section revealed that, caudally, the stacked rostrocaudal pattern did not overlap with the dorsoventral pattern. The former was contained in the Pax6 regime, whereas the latter fell fully inside the FB region. This resulted in five distinct regions in dorsoventral direction and nine in total.

### Overexpression simulations of morphogen secretions show high sensitivity of neural progenitor cell fate to morphogen levels

To investigate the influence of morphogen secretion levels, we simulated overexpression of SHH and WNT production. SSH oex did not affect the rostrocaudal pattern, but significantly changed the dorsoventral pattern (Figure 4 A). The Nkx2.2 domain was drastically dorsally enlarged compared to the wildtype (WT) and took up the whole ventral half of the neural tube. This led to a dorsal shift of the Olig2 domain, which, however, did not change in size. Consequently, the extent of the Pax6 domain was smaller. Through the dorsal shift of the pattern, the split of the dorsoventral domains occurred closer to the isthmic organiser and hence the extents of MB/HB + Olig2/Nkx2.2 domains were enlarged, leading to a decrease in size of the FB + Pax6/Olig2 domains.

**Figure 4.**
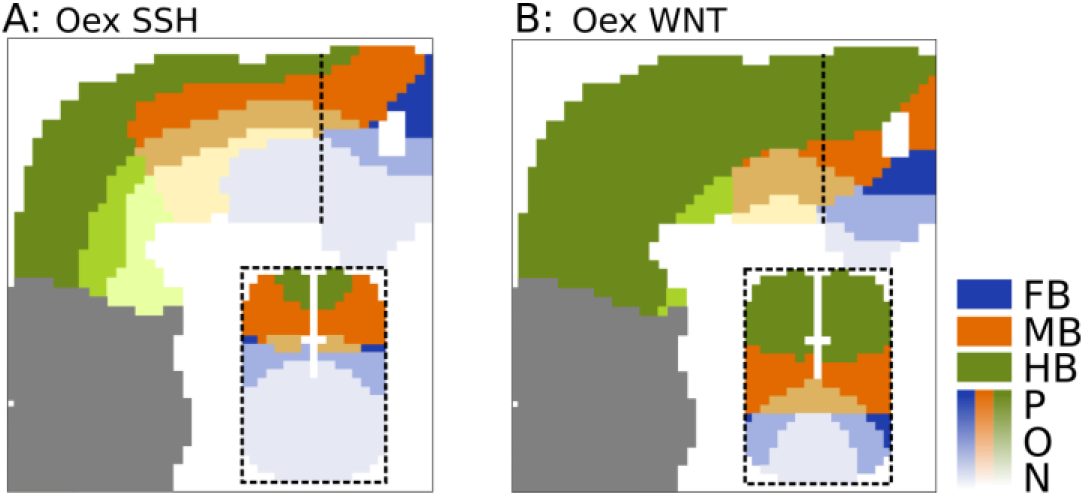
SHH and WNT overexpression simulations. A: Overexpression of SSH secretion. The rostrocaudal pattern was unaffected by the overexpression, whereas the Nkx2.2 domain was significantly larger than in the WT and the Pax6 domain was consequently smaller. B: Overexpression of WNT secretion. Predominantly, the rostrocaudal pattern was affected. The HB region was drastically enlarged, which led to smaller MB and FB regions. Moreover, the L-shape of WT HB and MB domains was lost. Greyed areas are only shown for orientation and are not considered for pattern establishment.

Overexpression of WNT had a small effect on dorsoventral patterning, decreasing the size of the Nkx2.2 domain and shifting the Olig2 and Pax6 domains ventrally. However, the effects of this overexpression on the rostrocaudal pattern were more striking (Figure 4 B). Most dominantly, the HB region was drastically enlarged, covering the whole caudal half of the neural tube and extending almost to the WT MB/FB boundary. This led to a ventral and caudal shift of the MB domain, which changed its characteristic L shape to a diagonal band and decreased its volume. Interestingly, through the dorsoventral stacking, the MB domain extended up to the caudal boundary of the neural tube. The FB domain was positioned in the ventral and rostral tip of the neural tube, covering less than half the volume of the WT FB domain. All this led to a more pronounced dorsoventral stacking of the rostrocaudal brain regions in the caudal half of the neural tube, which gave rise to a large MB + Olig2 domain, while the volume of the WT HB + Nkx2.2 domain was drastically reduced.

## Discussion

We presented a simple mathematical model of rostrocaudal neural tube patterning. We determined its topology in an unbiased and data-driven fashion. The resulting configuration was that of a tristable switch of mutually repressing brain region fate under the control of the WNT target GSK3. The unbiasedly discovered gene regulatory network has at its core genetic toggle switches consisting of cross-repressing transcription factors, which are network motifs that were shown to control patterning at tissue levels (39). The toggle switch motif has the capability of translating the continuous signal into an on-off behavior, in our case WNT gradient signal into e.g. FB off / MB on. Our multiple toggle switch model was able to recapitulate data from *in vitro* studies on hPSCs cultured at different WNT signalling levels (8) and exhibited similar dynamics as recent experiments done on a synthetic *in vitro* setup (26). The toggle switch motif has the capability of controlling the boundaries of gene expression domains (27,40). Thus, parameter perturbation or manipulation (e.g. via knockdown, overexpresion) of one of the factors can lead to changes in the resulting pattern. For our network model parameter sensitivity analysis and *in silico* overexpression and knockdown experiments suggested that the repression of HB fate, rather than the repression of FB fate or the promotion of MB fate, is the most promising strategy to increase efficiency and robustness of DA MB generation protocols by increasing the window of supplied GSK3 inhibitor CT that results in MB fate. It will be interesting to see if this result can be validated experimentally; one option could be to use synthetic neural tube patterning similar to the work recently published by Rifes et al. (26).

Combining the rostrocaudal model with a previously published model for dorsoventral neural tube patterning resulted in an integrated model controlled by the morphogens SHH and WNT. On the rostrocaudal side of this model, a buffer node U, which translates WNT signal to GSK3 inhibition, needed to be introduced to achieve stable steady state rostrocaudal patterning. This node corresponds most likely to one or more elements in the WNT pathway and the need for it is an example for the dynamic difference between signalling cascades and direct modulation of gene expression. It should be noted that the configuration of the rostrocaudal branch of the integrated network is very similar to the dorsoventral branch, with both branches having a morphogen level-regulated multiple toggle switch configuration at their core. This is very much in line with the idea that network motifs like feedback, mutual repression predominate parts of the modular structure of gene regulatory networks (39,41). To test the capabilities of the combined model, we simulated the patterning of the early human neural tube. For that, we set up a realistic three-dimensional model of the neural tube, which included the major morphogen secretion sites. The diffusion of the morphogens from these sites led to stable patterning of the tube into distinct regions. Interestingly, the bent geometry of the neural tube, combined with WNT secretion from the roof plate led to dorsoventral stacking of the rostrocaudal domains in the rostral half of the tube. It would be very interesting to find out if this is a shortcoming of the model or indeed a patterning mechanism. Lastly, we simulated the overexpression of morphogen secretion, which illustrated that cell fate and consequently neural tube patterning is sensitive to the abundance of morphogens seen by individual cells. In the WNT overexpression case, the model predicted an increase in HB fate, which is consistent with a study in Xenopus explants (42). Moreover, we find that the general sensitivity of neural progenitor cells to the abundance of WNT and SHH ligands predicted by the model to be in line with a recent study on hPSCs (43).

In summary, the presented model for rostrocaudal neural tube patterning can, despite its simplicity, recapitulate the main features of neural tube patterning and neural differentiation in dependence of WNT signalling and predict strategies of improving the efficiency of DA MB generation protocols. In the future, targeted experiments *in vitro* are required to test the predictions of this model. Moreover, our 3D computational model is a first step towards an integrated and conclusive view of neural tube patterning *in vivo*.

## Methods

### Clustering of gene expression data

For a gene with expression level *x* the normalisation was chosen such that 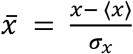 with the normalised gene expression 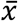, the mean gene expression 〈*x*〉 across the different CT levels and the corresponding standard deviation *σ*_*x*_.

The normalised gene expression data was clustered using hierarchical agglomerative clustering (30). As method for the computation of the cluster distance, the Nearest Point Algorithm was used, meaning that the minimal distance between two clusters was considered as their distance. The used metric for the determination of the distance was the Euclidean distance.

### Optimisation of model towards *in vitro* data

The parameters of the model were optimised using the experimental data. For the purpose of parameter optimisation, the expression of each brain region’s specific genes was normalised such that its maximum value was one via division of all values by the maximum value. During the optimisation, the network was initialised with FB = MB = HB = GSK3 = 1 for all used CT concentrations. The evolution of the gene expressions was computed iteratively, stabilising at *t* = 200. This steady state gene expression for a specific CT level was considered as the model output. The cost function *E_d_* measures the deviation of the model output *r* from the data *d* and is computed using

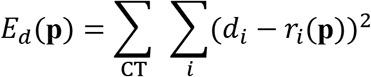

with the first sum going over the different CT concentrations, the second sum going over *i* ∈ {FB, MB, HB} and the parameter set ***p***. An optimal set of parameters is found by minimising *E_d_*(***p***) with respect to ***p*** using a L-BFGS-B algorithm (44). The parameter set used in this article is shown in Table 1.

**Table 1.** Model Parameters.

| $i$ | 1 | 2 | 3 | 4 | 5 | 6 | 7 | 8 |
| --- | --- | --- | --- | --- | --- | --- | --- | --- |
| $c_i$ | 0.015 | 8 | 1000 | 0.2 | 0.5 | 0.201 | 0.0048 | 0.205 |
| $n_i$ | 4 | 4 | 4 | 1 | 2 | 1 | 3 | 1 |
| $\delta_i$ | 0.169 | 0.169 | 0.171 | 0.171 | 0.1 | | | |
| $i$ | 9 | 10 | 11 | 12 | 13 | 14 | 15 | 16 |
| $c_i$ | 2 | 0.05 | 0.051 | 0.248 | 0.152 | 50 | 0.14 | 25 |
| $n_i$ | 3 | 1 | 3 | 1 | 1 | 2.5 | 1 | 4 |
| $\delta$ | $\delta_{WNT}$ | $\delta_{SHH}$ | $D_{WNT}$ | $D_{SHH}$ | $W_{critG}$ | h6 | $k_4$ | |
| 5.0 | 0.04 | 0.1 | 150.7 | 133.4 | 1.0 | 1.0 | 0.15 |  |

### Differential equations of *in vitro* gene regulatory network

The set of differential equations describing the *in vitro* model mathematically was constructed using the Hill formalism and resulted in

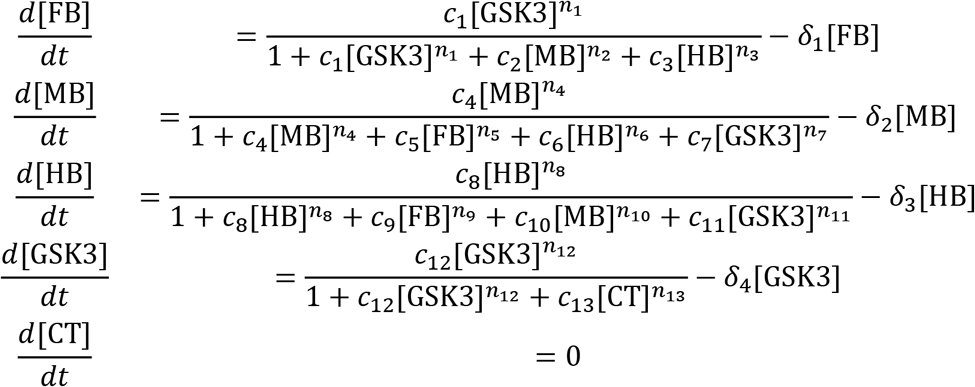

The set of differential equations defining the *in vivo* model are

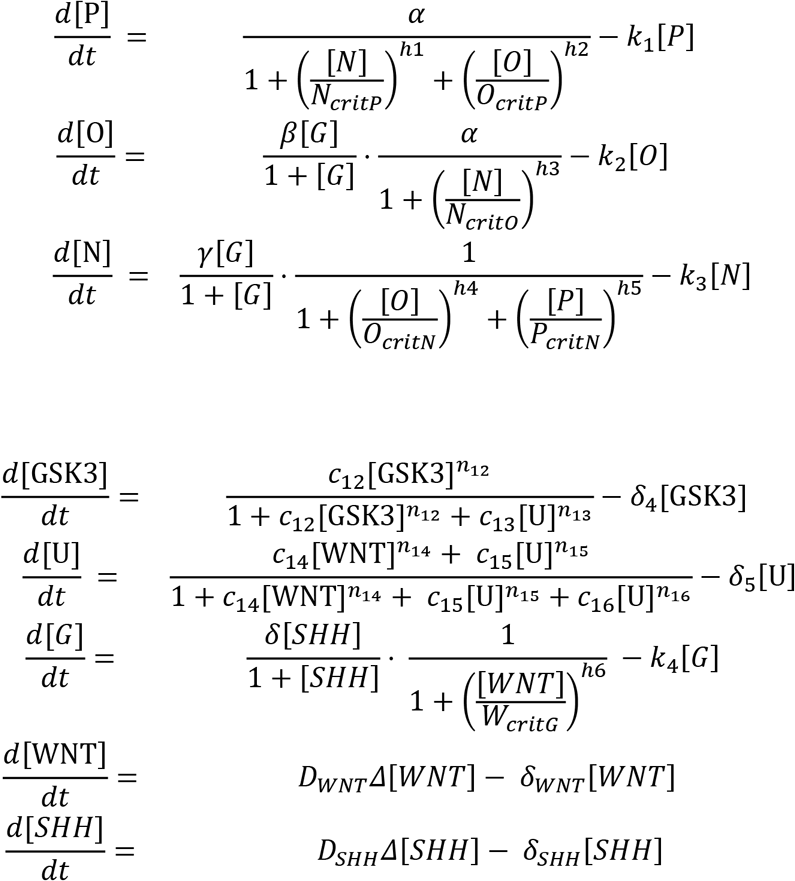

with the parameters in Table 1. The self-activation term of FB used during the model selection is not shown in the equations above, since it is not used after this step. The term was implemented in the ODE for FB expression following the Hill formalism, i.e. in the denominator for repression and in both numerator and denominator for activation.

The morphogen diffusion of the *in vivo* model was implemented using the discrete version of Fick’s law

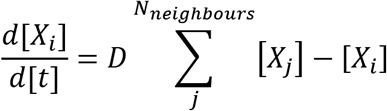

with the index of the current cell *i*, the diffusion constant *D*, and [*X*] being the concentration of either WNT or SHH. For each time step, the morphogen gradient pattern was updated prior to the intra-cellular network. The model reaches a steady state pattern, as shown in Figure 3 C.

### Model parameters and initial conditions

The model parameters are summarized in Table 1. Unlisted parameters of the dorsoventral model are listed in (27).

The FB self-interaction parameters were c_0_ = 0.0005 and *n*_0_ = 1.

For the *in vitro* simulations, the system was initialised with FB = MB = HB = GSK3 = 1. The 3D *in vivo* simulations were initialised with WNT = SHH = U = P = O = N = 0, FB = GSK3 = 1, MB = HB = 0.001, G = 3. Morphogen producing cells were held constant at WNT_prod_ = 2, SHH_prod_ = 1.

### Kd and oex implementation

The simulation of knockdown and overexpression *in vitro* was achieved by fixing the respective kd/oex node’s derivative to 0. The initial expression of the node then determined its behaviour, with kd being simulated by [ ]_0_ = 0 and oex [ ]_0_ = 1. The morphogen secretion overexpression *in vivo* was implemented by increasing the morphogen level of the secreting cells ten-fold. In all cases, the steady state expression pattern is considered the model output.

### Morphogen diffusivity

The morphogens’ diffusion coefficients *D_WNT_*, *D_SHH_* were estimated following a method proposed by He and Niemeyer based on the radius of gyration *R_G_* of the protein (45). *R_G_* was estimated according to (46) for physiological conditions:

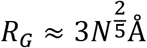

with N being the protein chain length (number of amino acids). Inserting the protein lengths *N_WNT_* = 370 and *N_SHH_* = 462 (47) yields

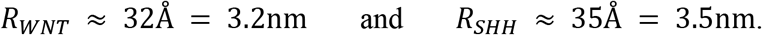

Subsequently, we applied the estimate

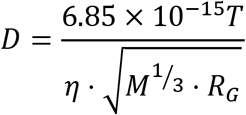

with *D* in units of m^2^·s^−1^, temperature *T* = 310 *K*, viscosity *η* = 0.75 cP (48) and the molecular Mass *M* (*M*_*WNT*_ = 40.98*kDa*, *M*_*SHH*_ = 49.61*kDa*) in kg ·kmol^−1^, yielding

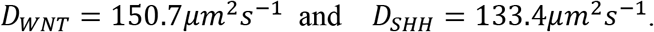

### Computational methods

The *in vitro* model equations were solved numerically using a fourth order Runge-Kutta algorithm with temporal step size *h* = 2. The equations of the *in vivo* model were solved using the explicit Euler method.

All computational work was performed in Python 3.7 with the extensions NumPy (49), SciPy (50) and Matplotlib (51). The neural tube model and the sources of morphogen secretion were set up in MagicVoxel, an open source voxel editor software. Cells were approximated as cubes with side length 10μm with the morphogen secretions sites positioned as illustrated in Figure 3 B.

## Acknowledgements

The authors would like to thank Julia Deichmann for discussions at various stages of the project and for critical input on the manuscript.

## References

1. Chinta SJ, Andersen JK. Dopaminergic neurons. International Journal of Biochemistry and Cell Biology. 2005.

2. Takagi Y, Takahashi J, Saiki H, Morizane A, Hayashi T, Kishi Y, et al. Dopaminergic neurons generated from monkey embryonic stem cells function in a Parkinson primate model. J Clin Invest. 2005;

3. Lehnen D, Barral S, Cardoso T, Grealish S, Heuer A, Smiyakin A, et al. IAP-Based Cell Sorting Results in Homogeneous Transplantable Dopaminergic Precursor Cells Derived from Human Pluripotent Stem Cells. Stem Cell Reports. 2017;

4. Kefalopoulou Z, Politis M, Piccini P, Mencacci N, Bhatia K, Jahanshahi M, et al. Long-term clinical outcome of fetal cell transplantation for parkinson disease: Two case reports. JAMA Neurol. 2014;

5. Barker RA, Drouin-Ouellet J, Parmar M. Cell-based therapies for Parkinson disease-past insights and future potential. Nature Reviews Neurology. 2015.

6. Lindvall O. Dopaminergic neurons for Parkinson’s therapy. Nature Biotechnology. 2012.

7. Grealish S, Diguet E, Kirkeby A, Mattsson B, Heuer A, Bramoulle Y, et al. Human ESC-derived dopamine neurons show similar preclinical efficacy and potency to fetal neurons when grafted in a rat model of Parkinson’s disease. Cell Stem Cell. 2014;

8. Kirkeby A, Parmar M. Building authentic midbrain dopaminergic neurons from stem cells - Lessons from development. Transl Neurosci. 2012;3(4):314–9.

9. Kriks S, Shim J-W, Piao J, Ganat YM, Wakeman DR, Xie Z, et al. Dopamine neurons derived from human ES cells efficiently engraft in animal models of Parkinson’s disease. Nature [Internet]. 2011; Available from: http://www.nature.com/doifinder/10.1038/nature10648

10. Steinbeck JA, Choi SJ, Mrejeru A, Ganat Y, Deisseroth K, Sulzer D, et al. Optogenetics enables functional analysis of human embryonic stem cell-derived grafts in a Parkinson’s disease model. Nat Biotechnol. 2015;

11. Kirkeby A, Grealish S, Wolf DA, Nelander J, Wood J, Lundblad M, et al. Generation of Regionally Specified Neural Progenitors and Functional Neurons from Human Embryonic Stem Cells under Defined Conditions. Cell Rep [Internet]. 2012 Jun;1(6):703–14. Available from: http://linkinghub.elsevier.com/retrieve/pii/S2211124712001222

12. Nolbrant S, Heuer A, Parmar M, Kirkeby A. Generation of high-purity human ventral midbrain dopaminergic progenitors for in vitro maturation and intracerebral transplantation. Nat Protoc [Internet]. 2017;12(9):1962–79. Available from: http://www.nature.com/doifinder/10.1038/nprot.2017.078

13. González H, Contreras F, Prado C, Elgueta D, Franz D, Bernales S, et al. Dopamine Receptor D3 Expressed on CD4 + T Cells Favors Neurodegeneration of Dopaminergic Neurons during Parkinson’s Disease. J Immunol. 2013;

14. Adil MM, Rodrigues GMC, Kulkarni RU, Rao AT, Chernavsky NE, Miller EW, et al. Efficient generation of hPSC-derived midbrain dopaminergic neurons in a fully defined, scalable, 3D biomaterial platform. Sci Rep. 2017;

15. Cho MS, Hwang DY, Kim DW. Efficient derivation of functional dopaminergic neurons from human embryonic stem cells on a large scale. Nat Protoc. 2008;

16. Kim JH, Auerbach JM, Rodríguez-Gómez JA, Velasco I, Gavin D, Lumelsky N, et al. Dopamine neurons derived from embryonic stem cells function in an animal model of Parkinson’s disease. Nature. 2002;

17. Gasser RF. Atlas of human embryos. Hagerstown: Harper & Row; 1975. ix, 317 p. :

18. Wilson L, Maden M. The mechanisms of dorsoventral patterning in the vertebrate neural tube. Developmental Biology. 2005.

19. Ulloa F, Martí E. Wnt won the war: Antagonistic role of Wnt over Shh controls dorso-ventral patterning of the vertebrate neural tube. Developmental Dynamics. 2010.

20. Carlson BM. Human Embryology and Developmental Biology: Fifth Edition. Human Embryology and Developmental Biology: Fifth Edition. 2013.

21. Peter IS, Faure E, Davidson EH. Predictive computation of genomic logic processing functions in embryonic development. Proc Natl Acad Sci U S A [Internet]. 2012 Oct 9 [cited 2020 Aug 1];109(41):16434–42. Available from: https://www.pnas.org/content/109/41/16434

22. Peter IS, Davidson EH. Genomic Control Process: Development and Evolution. Genomic Control Process: Development and Evolution. Elsevier Inc.; 2015. 1–448 p.

23. Sagner A, Briscoe J. Morphogen interpretation: concentration, time, competence, and signaling dynamics. Wiley Interdiscip Rev Dev Biol [Internet]. 2017 Jul 1 [cited 2020 Aug 1];6(4):e271. Available from: http://doi.wiley.com/10.1002/wdev.271

24. Briscoe J. Understanding Pattern Formation in Embryos: Experiment, Theory, and Simulation. J Comput Biol [Internet]. 2019 Jul 1 [cited 2020 Aug 1];26(7):696–702. Available from: https://www.liebertpub.com/doi/10.1089/cmb.2019.0090

25. Tao Y, Zhang SC. Neural Subtype Specification from Human Pluripotent Stem Cells. Cell Stem Cell. 2016.

26. Rifes P, Isaksson M, Rathore GS, Aldrin-Kirk P, Møller OK, Barzaghi G, et al. Modeling neural tube development by differentiation of human embryonic stem cells in a microfluidic WNT gradient. Nat Biotechnol [Internet]. 2020 May 25 [cited 2020 Jun 25];1–9. Available from: https://www.nature.com/articles/s41587-020-0525-0

27. Balaskas N, Ribeiro A, Panovska J, Dessaud E, Sasai N, Page KM, et al. Gene regulatory logic for reading the sonic hedgehog signaling gradient in the vertebrate neural tube. Cell. 2012;

28. Monzel AS, Smits LM, Hemmer K, Hachi S, Moreno EL, van Wuellen T, et al. Derivation of Human Midbrain-Specific Organoids from Neuroepithelial Stem Cells. Stem Cell Reports. 2017;

29. Smits LM, Reinhardt L, Reinhardt P, Glatza M, Monzel AS, Stanslowsky N, et al. Modeling Parkinson’s disease in midbrain-like organoids. npj Park Dis. 2019;

30. Müllner D. Modern hierarchical, agglomerative clustering algorithms. arXiv Prepr. 2011;(1973):1–29.

31. Kobayashi D, Kobayashi M, Matsumoto K, Ogura T, Nakafuku M, Shimamura K. Early subdivisions in the neural plate define distinct competence for inductive signals. Development. 2002;

32. Xuan S, Baptista CA, Balas G, Tao W, Soares VC, Lai E. Winged helix transcription factor BF-1 is essential for the development of the cerebral hemispheres. Neuron. 1995;

33. Toresson H, Martinez-Barbera JP, Bardsley A, Caubit X, Krauss S. Conservation of BF-1 expression in amphioxus and zebrafish suggests evolutionary ancestry of anterior cell types that contribute to the vertebrate telencephalon. Dev Genes Evol. 1998;

34. Alves dos Santos MTM, Smidt MP. En1 and Wnt signaling in midbrain dopaminergic neuronal development. Neural Development. 2011.

35. Kouwenhoven WM, Von Oerthel L, Smidt MP. Pitx3 and En1 determine the size and molecular programming of the dopaminergic neuronal pool. PLoS One. 2017;

36. Barrow JR, Stadler HS, Capecchi MR. Roles of Hoxa1 and Hoxa2 in patterning the early hindbrain of the mouse. Development. 2000;

37. Davenne M, Maconochie MK, Neun R, Pattyn A, Chambon P, Krumlauf R, et al. Hoxa2 and Hoxb2 control dorsoventral patterns of neuronal development in the rostral hindbrain. Neuron. 1999;

38. De Bakker BS, De Jong KH, Hagoort J, De Bree K, Besselink CT, De Kanter FEC, et al. An interactive three-dimensional digital atlas and quantitative database of human development. Science (80-). 2016;

39. Davidson EH. Emerging properties of animal gene regulatory networks [Internet]. Vol. 468, Nature. Nature Publishing Group; 2010 [cited 2020 Aug 1]. p. 911–20. Available from: http://sugp.caltech.edu/endomes/.

40. Briscoe J, Pierani A, Jessell TM, Ericson J. A homeodomain protein code specifies progenitor cell identity and neuronal fate in the ventral neural tube. Cell [Internet]. 2000 May 12 [cited 2020 Aug 1];101(4):435–45. Available from: http://www.cell.com/article/S0092867400808533/fulltext

41. Alon U. An Introduction to Systems Biology: Design Principles of Biological Circuits (Chapman and Hall/CRC Mathematical and Computational Biology). Star. 2006.

42. Itoh K, Sokol SY. Graded amounts of Xenopus dishevelled specify discrete anteroposterior cell fates in prospective ectoderm. Mech Dev. 1997;

43. Strano A, Tuck E, Stubbs VE, Livesey FJ. Variable Outcomes in Neural Differentiation of Human PSCs Arise from Intrinsic Differences in Developmental Signaling Pathways. Cell Rep. 2020;

44. Byrd RH, Lu P, Nocedal J, Zhu C. A Limited Memory Algorithm for Bound Constrained Optimization. SIAM J Sci Comput [Internet]. 1995;16(5):1190–208. Available from: http://epubs.siam.org/doi/10.1137/0916069

45. He L, Niemeyer B. A novel correlation for protein diffusion coefficients based on molecular weight and radius of gyration. Biotechnol Prog. 2003;19(2):544–8.

46. Hong L, Lei J. Scaling law for the radius of gyration of proteins and its dependence on hydrophobicity. J Polym Sci Part B Polym Phys. 2009;

47. UniProt Consortium T. UniProt: the universal protein knowledgebase. Nucleic Acids Res. 2018;

48. Puchkov E. Intracellular viscosity: Methods of measurement and role in metabolism. Biochem Suppl Ser A Membr Cell Biol [Internet]. 2013;7(4):270–9. Available from: http://link.springer.com/10.1134/S1990747813050140

49. Van Der Walt S, Colbert SC, Varoquaux G. The NumPy array: A structure for efficient numerical computation. Comput Sci Eng. 2011;

50. Virtanen P, Gommers R, Oliphant TE, Haberland M, Reddy T, Cournapeau D, et al. SciPy 1.0: fundamental algorithms for scientific computing in Python. Nat Methods. 2020;

51. Hunter JD. Matplotlib: A 2D graphics environment. Comput Sci Eng. 2007;

